# Adverse prognosis of GM-CSF expression in human cutaneous melanoma

**DOI:** 10.1101/2025.05.30.656980

**Authors:** Elena García-Martínez, Alicia Nieto-Valle, Celia Barrio-Alonso, Baltasar López-Navarro, José Antonio Avilés-Izquierdo, Verónica Parra-Blanco, Alba Gutiérrez-Seijo, Paloma Sánchez-Mateos, Rafael Samaniego

**Affiliations:** Unidad de Microscopía Confocal, Instituto de Investigación Sanitaria Gregorio Marañón, Madrid, Spain; Laboratorio de Inmuno-oncología, Instituto de Investigación Sanitaria Gregorio Marañón, Madrid, Spain; Departamento de Inmunología, Universidad Complutense de Madrid, Madrid, Spain; Servicio de Dermatología, Instituto de Investigación Sanitaria Gregorio Marañón, Madrid, Spain; Servicio de Anatomía Patológica, Instituto de Investigación Sanitaria Gregorio Marañón, Madrid, Spain

## Abstract

Tumor-associated macrophages (TAMs) represent a major immune population within the tumor microenvironment, influencing cancer progression and immune responses. Our group previously identified a subset of pro-inflammatory TAMs associated with poor prognosis in human melanoma. GM-CSF, a myeloid-priming cytokine, exhibits context-dependent effects on tumor growth and, despite its clinical use, its role in human melanoma remains undefined. In this study, we demonstrate that GM-CSF is significantly enriched in primary cutaneous melanoma samples from patients who subsequently developed metastasis, compared with non-metastasizing ones. By quantifying GM-CSF expression in TAMs and tumor cells (TCs), we found that high levels, in both TAMs and TCs, correlated with reduced disease-free and overall survival (p< 0.0001). Although GM-CSF receptor subunits were present in TCs and TAMs, their expression did not correlate with clinical outcomes. To explore the pro-metastatic role of GM-CSF, we performed in vitro assays and found that it activates non-canonical signaling pathways, such as STAT3 and PI3K/AKT, and promotes melanoma cell invasion. Consistently, its administration enhanced lung colonization by melanoma cells in a murine model, an effect reversed by CD116 receptor blockade. Furthermore, GM-CSF–primed macrophages secreted higher levels of inflammatory cytokines upon interaction with melanoma cells than those unprimed or primed with the counterpart cytokine M-CSF. Reciprocally, RNA-seq analyses revealed a broader transcriptional reprogramming in melanoma cells exposed to GM-CSF-primed macrophages, which displayed enhanced expression of inflammatory-response genes, suggesting a feedforward loop. Altogether, our findings highlight a potentially pro-tumorigenic GM-CSF-driven paracrine axis in patients with poor-prognosis melanoma, supporting therapeutic strategies aimed at disrupting this signaling network.

## INTRODUCTION

Tumor-associated macrophages (TAMs) are typically the most abundant leukocytes within the tumor microenvironment, where they play a central role in immunoregulation and tumor progression. Our group has shown that, in melanoma patients, TAMs exhibiting a cytokine-producing pro-inflammatory phenotype, consistent with the recently defined Inflam-TAM subset (1,2), are associated with poor prognosis and increased occurrence of distant metastasis (3). Granulocyte-macrophage colony-stimulating factor (GM-CSF), along with macrophage colony-stimulating factor (M-CSF), are well-established growth factors that play critical roles in promoting the recruitment, survival, and differentiation of monocytes into macrophages. These factors induce distinct and often opposing polarization profiles in macrophages, particularly in the context of inflammation and immune responses (4). GM-CSF is usually associated with pro-inflammatory responses, and the transcriptional response of monocytes to GM-CSF seems to be much more extensive than that to M-CSF, leading to an enhanced polarization and a more terminally differentiated state in macrophages (5).

GM-CSF is secreted by various immune and non-immune cells, including cancer cells, in response to inflammatory or damage-associated stimuli. The regulation of its functions is partly dictated by the biology of its receptor (GM-CSFR), which consists of an α subunit (CD116) responsible for ligand binding and a β subunit (CD131) that mediates intracellular activation through JAK2. This engagement can trigger the canonical JAK2/STAT5 pathway and can also activate other signaling cascades, such as Ras/MAPK and PI3K/AKT. Both JAK2/STAT5 and MAPK pathways have been linked with cellular activation, inflammation, and proliferation, while PI3K/AKT signaling seems more associated with enhanced cell survival and inhibition of apoptosis (6,7). Additionally, activation of the NF-κB pathway has been reported through direct interaction of IKKβ with the CD116 subunit, contributing to survival, proliferative and tolerogenic responses (8); and other transcription factors associated with JAK2, such as STAT3, have also been described as part of GM-CSF-mediated signaling, promoting the expression of molecules involved in immunosuppressive pathways (9,10). Several factors may influence the signaling outcomes and functional effects of the GM-CSF/GM-CSFR axis, including cellular type, ligand concentration, receptor expression levels, and its structural organization, which can exist in either tetrameric or dodecameric forms (8,11). Evidence suggests that GM-CSF, rather than acting as a definitive polarizing signal, primarily functions as a priming stimulus, with the final immune response being shaped by additional factors present in the microenvironment (12,13).

Remarkably, despite its conventional association with a pro-inflammatory phenotype, GM-CSF also exhibits a dual role, displaying immunoregulatory properties under certain circumstances (6). Consistently, GM-CSF has been implicated in both pro-tumoral and anti-tumoral roles within the tumor microenvironment. The expression of GM-CSF has been studied across various tumor types, with distinct associations to patient prognosis (11). In addition, numerous in vitro and murine studies have examined the role of GM-CSF, demonstrating its impact on tumor development and progression through both immune-dependent mechanisms -such as modulation of myeloid cell recruitment and activation- and immune-independent pathways involving direct effects on tumor cells (14).

Specifically in melanoma, murine studies using the B16 cell line demonstrated that GM-CSF increased tumor growth and angiogenesis, and that this effect was dependent on the interaction and recruitment of bone marrow-derived cells (15). By contrast, other studies using murine xenograft models with the human melanoma cell line A375 demonstrated that treatment with GM-CSF in induced hypoxic conditions, reduced tumor growth by enhancing sVEGFR-1 production from TAMs (16). Such conflicting results have also been highlighted by other authors, who, using xenograft mouse models with distinct human melanoma cell lines, reported heterogeneous inter-tumor responses to GM-CSF (17). However, to date, no studies have specifically addressed the expression of GM-CSF/GM-CSFR in human melanoma, despite its inclusion in therapeutic strategies for patients with this disease (18).

In this study, we examined the expression of GM-CSF and its receptor in primary melanoma tumor samples from our patient cohort, exploring their potential correlation with metastasis development. Additionally, we investigated the effects of GM-CSF on tumor cells (TCs) and its role in shaping tumor-associated macrophages profiles within the melanoma microenvironment.

## MATERIAL AND METHODS

### Study cohort and selection criteria

Patient samples were collected following the approval of the Hospital General Universitario Gregorio Marañón ethics committee. A formalin-fixed and paraffin-embedded (FFPE) primary cutaneous melanoma cohort of 80 patients was used, with a >2 mm Breslow thickness, Pathological American Joint Committee on Cancer staging II–IV, and a median follow-up of 81 months, as previously described (3). Six patients at stage IV were excluded from disease-free survival (DFS) analysis, but not from overall survival (OS).

### Fluorescence confocal microscopy

FFPE samples were deparaffinized, rehydrated, and unmasked by steaming in 10 mM sodium citrate buffer pH 9.0 (Dako, Glostrup, Denmark) for 6 min. Slides were blocked with 5 μg/mL human immunoglobulins solved in blocking serum-free medium (Dako) for 30 min and then sequentially incubated with 5–10 μg/mL primary antibodies (supplementary Table S1) and appropriate fluorescent secondary antibodies (Jackson Immunoresearch, West Grove, PA, USA), as previously described (3). Samples were imaged with a SPE confocal microscope using the glycerol-immersion ACS APO x20/NA 0.60 objective (Leica Microsystems, Wetzlar, Germany). Single-cell quantification was performed at 3-5 20x fields, and Mean Fluorescence Intensity (MFI) of proteins obtained at manually depicted TCs or at segmented CD68^+^ TAMs using the ‘analyze particle’ plugin of ImageJ2 software, as previously described (19). For in vivo lung colonization assays, NG2^+^ melanoma micrometastases were microscopically measured at multiple histologic frozen sections (20).

### Monocyte Isolation and Cell Culture

Peripheral blood mononuclear cells (PBMCs) were isolated from buffy-coats from healthy donors over a ficoll gradient (Lymphocytes Isolation Solution, Rafer). Monocytes were purified by magnetic cell sorting using anti-CD14 tagged microbeads (Miltenyi Biotech, Bergisch Gladbach, Germany). For in vitro priming of macrophages, monocytes were cultured at 0.5 × 10^6^/mL for 7 days containing recombinant human GM-CSF (10 ng/mL, Immunotools, Friesoythe, Germany) or recombinant human M-CSF (10 ng/mL, Immunotools) to generate GM-MP or M-MP macrophages, respectively. Cytokines were added every two days. All cells, including the melanoma cell lines BLM, A375, and Skmel-103 (21) were cultured in RMPI-1640 medium (Gibco, Waltham, MA, USA) supplemented with 10% fetal calf serum (FCS, Sigma, Burlington, MA, USA).

### In vitro mRNA measurements

Melanoma cells were co-cultured with monocytes or primed macrophages at 1:2 ratio (melanoma:myeloid). Conditioned media were collected after 48h for CCL20, Activin A, IL-1β (R&D Systems, Minneapolis) and GM-CSF (Biolegend, San Diego, California, USA) secretion assessment. For transcriptomic analysis, TCs cells and monocytes/macrophages were processed after 24h co-culture, separating TCs from myeloid counterpart with anti-CD14 microbeads. Oligonucleotides (Supplementary Table S1) were designed according to the Roche software for real time quantitative PCR (qPCR) analysis (Roche Diagnostics, Basel, Switzerland), as previously described (21). Assays were made in triplicate and normalized to TBP expression (ΔΔCT method). Total RNA was also processed and sequenced at BGI (https://www.bgitechsolutions.com) using the DNBseq-G400 platform. DEGs were assessed by using DESeq2 algorithm with parameters log2 (fold change) >2 and adjusted p-value <0.05. DEGs were classified using the ‘enricher’ function of *clusterProfiler* R package (22) according to the hallmark gene sets of the Molecular Signature Database (MSigDB, https://www.gsea-msigdb.org/gsea/msigdb). Single sample Gene Set Enrichment Analysis (ssGSEA) was performed using the *GSVA* R package for several cytokine production gene sets from the “GO: Biological Process” subcategory of the MSigDB. Data were deposited in NCBI’s Gene Expression Omnibus and are accessible through GEO Series accession numbers GSE171277, GSE242674 and GSE280027.

### In vitro protein measurements

For protein expression analysis of GM-CSFR subunits, whole-cell lysates were subjected to western blotting with the indicated antibodies (supplementary Table S1). For signaling kinetics, human melanoma cell lines (BLM, A375, and Skmel-103) were seeded at a density of 1×10^6^ cells per well in 6-well plates and incubated for 3 hours in complete medium to allow cell adhesion. Then, adhered melanoma cells or monocytes (2.5×10^6^ per well) were starved in serum-free medium for 1 hour before supplementation with distinct human GM-CSF concentrations for 15 and 30 minutes. Cell lysates were directly prepared using 1x Laemmli sample buffer.

### Invasion and survival assays

BLM and Skmel-103 spheroids were produced by seeding 10^4^ cells per well in a 96-well U-bottom plate (ThermoFisher Scientific, Waltham, MA, USA) for 5–7 days, embedded in 2 mg/mL collagen type-I (StemCell Technologies, Vancouver, Canada) gels and allowed to invade for 72 hours in the presence of serum-deprived media ±GM-CSF or M-CSF, and ±2 μg/mL of neutralizing antibodies for human CD116 or isotype-control antibodies (supplementary Table S1). TCs and monocytes/macrophages were first co-cultured for 48 hours in 10% FCS medium, and then media were replaced by serum-free medium containing bovine serum albumin (BSA, 1%) for 24 additional hours to obtain the CM used in invasion assays. Quantifications were performed as previously described (20). For cell viability assays, melanoma cells were cultured in serum-deprived media with or without GM-CSF for 5 days. Media were then removed, and cells fixed with 4% formaldehyde for colorimetric crystal violet assay (Thermofisher Scientific).

### Melanoma xenograft models

NOD/Scid/Il2rg^−/−^ mice (NSG, The Jackson Laboratory, Bar Harbor, ME) were maintained under specific pathogen-free conditions. For lung colonization experiments, male mice were used at the age of 6-8 weeks (∼23 g). At 15 minute intervals, mice were consecutively injected with intraperitoneal 100 μl PBS ±25 μg neutralizing or control antibodies (supplementary Table S1); subcutaneous 100 μl PBS ±5 μg human GM-CSF, and intravenous tail vein injection of 0.5 × 10^6^ melanoma cells suspended in 150 μl PBS ±500 ng/mL hGM-CSF and ±2 μg/mL antibodies. Two additional daily doses of intraperitoneal antibodies and subcutaneous cytokines were administered, and at day 6 (BLM) or 10 (A375) lungs were extracted for quantitative histological analyses. Note that human GM-CSF only binds to human CD116 (23). Procedures were approved by the IiSGM animal care/use and Comunidad de Madrid committees (Proex: 296.4/22).

### Statistical analyses

Kaplan–Meier curves were used to analyze the correlation with patient DFS and OS using Youden’s index to determine where the cutoff point was equally specific and sensitive. The Cox regression method (univariate and multivariate) was used to identify independent prognostic variables and Mann–Whitney tests to evaluate the association with clinicopathological features. Paired and unpaired t-test and log-rank analyses were also used in this study (GraphPad software, San Diego, CA, USA), as indicated; p < 0.05 was considered statistically significant.

## RESULTS

### Expression of M-CSF, GM-CSF and GM-CSFR subunits in primary melanoma tissues and association with patient survival

To contextualize the potential roles of M-CSF and GM-CSF cytokines in cutaneous melanoma, we first analyzed their expression by multiplexed immunofluorescence, at single cell resolution, in melanoma tissues. We analyzed an 80-patient cohort of stage II-IV paraffin-embedded cutaneous melanomas with clinically annotated survival data, differentiating whether patients developed distant metastasis or not for up to 10 years following tumor resection (Fig. 1A). Notably, whereas M-CSF was generally expressed by tumor cells, GM-CSF expression was significantly enriched in both TAMs and TCs of metastasizing primary tumors, compared with non-metastasizing ones (Fig. 1B). Remarkably, GM-CSFR subunits CD116 and CD131 were expressed by melanoma cells in most primary tumors, regardless of whether metastasis eventually occurred (Fig. 1A and 1B). To confirm the prognostic value of GM-CSF in our cohort of melanomas we stratified its expression in TAMs and TCs and performed Kaplan-Meier analyses. High expression by both cell types was significantly associated with poor disease-free (Log rank, p< 0.0001) and overall (p< 0.007) survival (Fig. 1C and 1D and supplementary Table S2). Moreover, Cox regression multivariate analyses determined that GM-CSF expression by melanoma cells was a prognostic marker for disease-free (HR= 7.5, p< 0.001) and overall survival (HR= 6.9, p< 0.001), independent of age, sex, Breslow level and staging, as well as TAM GM-CSF expression for DFS (Table 1). As expected, GM-CSF levels were enriched in samples where pro-inflammatory ‘cytokine-producing’ TAMs were majority (Fig. 1E), according to our previous cohort classification (3). Altogether, our data determine that the tumor microenvironment of metastasizing primary melanomas is enriched in GM-CSF, which might exert a pro-metastatic function acting on macrophages and/or tumor cells themselves, as both cell types express the cytokine receptor.

**Table 1.**
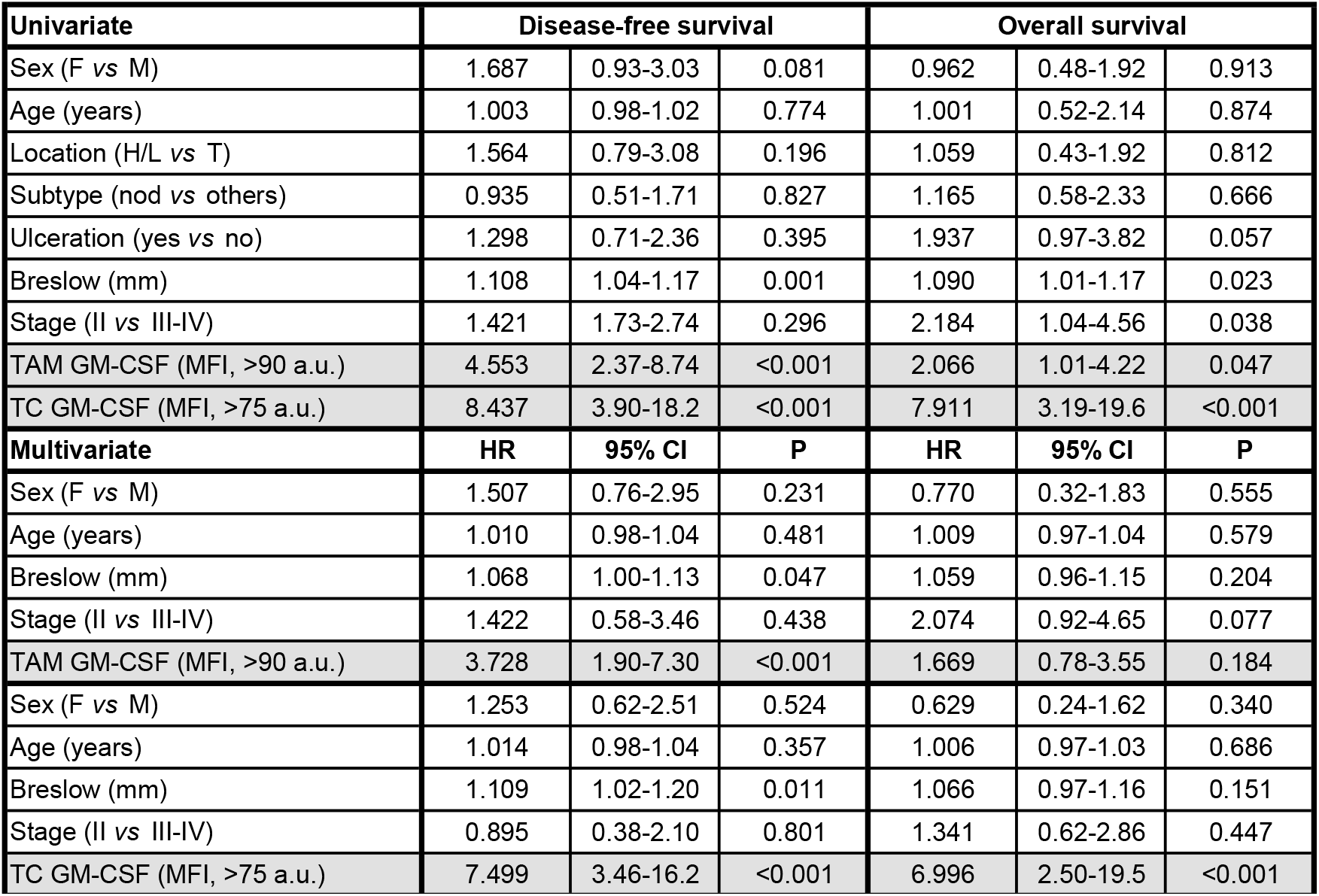
Univariate and multivariate Cox regression analyses for 10-year disease-free (DFS) and overall (OS) survival.

| Univariate | Disease-free survival |  |  | Overall survival |  |  |
| --- | --- | --- | --- | --- | --- | --- |
| Sex (F vs M) | 1.687 | 0.93-3.03 | 0.081 | 0.962 | 0.48-1.92 | 0.913 |
| Age (years) | 1.003 | 0.98-1.02 | 0.774 | 1.001 | 0.52-2.14 | 0.874 |
| Location (H/L vs T) | 1.564 | 0.79-3.08 | 0.196 | 1.059 | 0.43-1.92 | 0.812 |
| Subtype (nod vs others) | 0.935 | 0.51-1.71 | 0.827 | 1.165 | 0.58-2.33 | 0.666 |
| Ulceration (yes vs no) | 1.298 | 0.71-2.36 | 0.395 | 1.937 | 0.97-3.82 | 0.057 |
| Breslow (mm) | 1.108 | 1.04-1.17 | 0.001 | 1.090 | 1.01-1.17 | 0.023 |
| Stage (II vs III-IV) | 1.421 | 1.73-2.74 | 0.296 | 2.184 | 1.04-4.56 | 0.038 |
| TAM GM-CSF (MFI, >90 a.u.) | 4.553 | 2.37-8.74 | <0.001 | 2.066 | 1.01-4.22 | 0.047 |
| TC GM-CSF (MFI, >75 a.u.) | 8.437 | 3.90-18.2 | <0.001 | 7.911 | 3.19-19.6 | <0.001 |
| Multivariate | HR | 95% CI | P | HR | 95% CI | P |
| Sex (F vs M) | 1.507 | 0.76-2.95 | 0.231 | 0.770 | 0.32-1.83 | 0.555 |
| Age (years) | 1.010 | 0.98-1.04 | 0.481 | 1.009 | 0.97-1.04 | 0.579 |
| Breslow (mm) | 1.068 | 1.00-1.13 | 0.047 | 1.059 | 0.96-1.15 | 0.204 |
| Stage (II vs III-IV) | 1.422 | 0.58-3.46 | 0.438 | 2.074 | 0.92-4.65 | 0.077 |
| TAM GM-CSF (MFI, >90 a.u.) | 3.728 | 1.90-7.30 | <0.001 | 1.669 | 0.78-3.55 | 0.184 |
| Sex (F vs M) | 1.253 | 0.62-2.51 | 0.524 | 0.629 | 0.24-1.62 | 0.340 |
| Age (years) | 1.014 | 0.98-1.04 | 0.357 | 1.006 | 0.97-1.03 | 0.686 |
| Breslow (mm) | 1.109 | 1.02-1.20 | 0.011 | 1.066 | 0.97-1.16 | 0.151 |
| Stage (II vs III-IV) | 0.895 | 0.38-2.10 | 0.801 | 1.341 | 0.62-2.86 | 0.447 |
| TC GM-CSF (MFI, >75 a.u.) | 7.499 | 3.46-16.2 | <0.001 | 6.996 | 2.50-19.5 | <0.001 |

**Figure 1.**
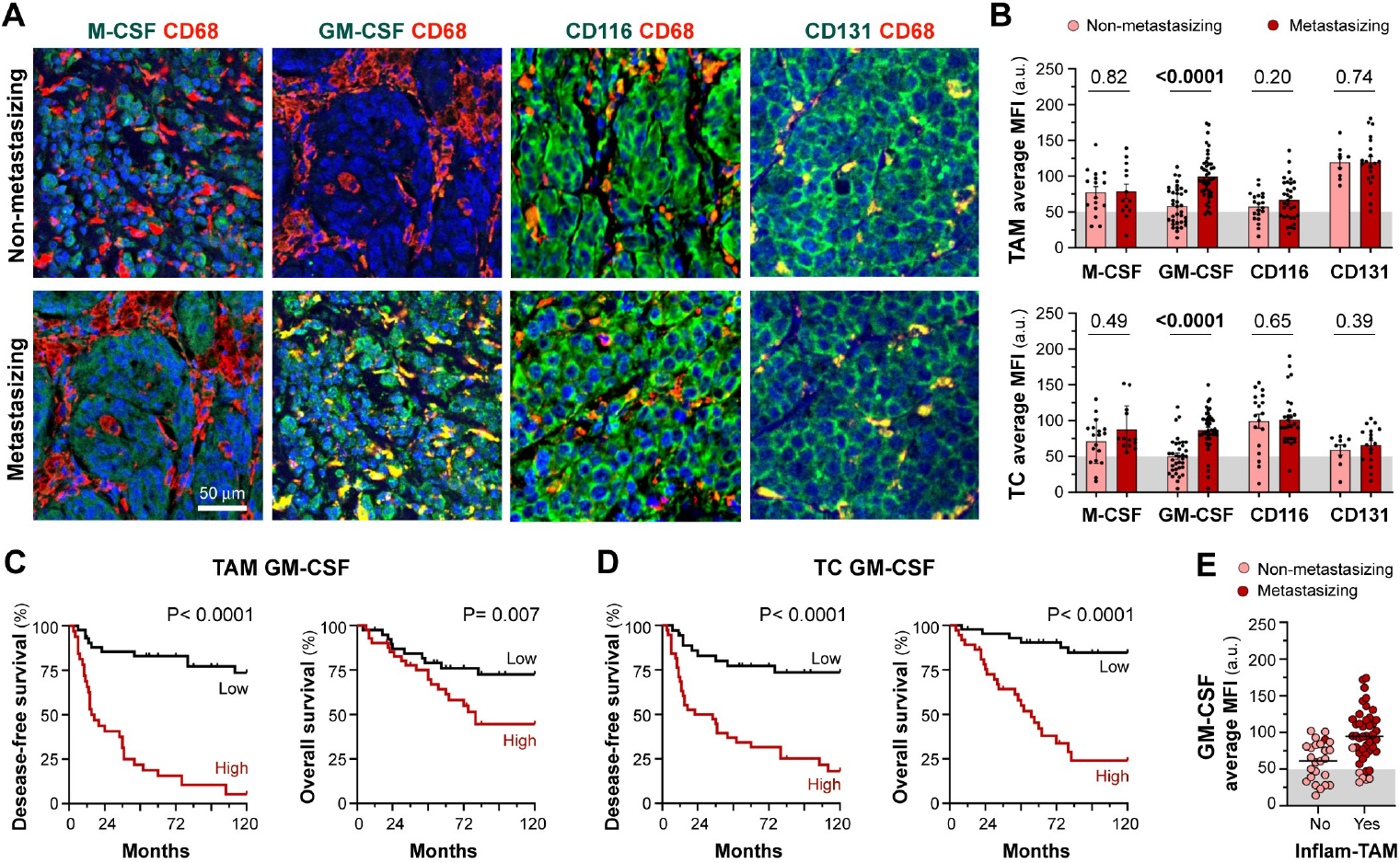
Confocal microscopy analysis of M-CSF, GM-CSF and GM-CSFR expression in human primary melanomas. **A**, representative FFPE sections of non-metastasizing and metastasizing primary melanoma samples co-stained for the pan-macrophage marker CD68 (red) and the proteins of interest (green), as indicated. **B**, single-cell quantification of M-CSF, GM-CSF, CD116 and CD131 in CD68^+^ TAMs and TCs. Plots show the average MFIs (arbitrary units, a.u.) of analyzed proteins for each tumor sample. Mann–Whitney *p* values, are shown. **C, D**, disease-free and overall survival 10-year Kaplan–Meier curves for GM-CSF expression in TAMs (**C**) and TCs (**D**). Log-rank *p* values, are shown. **E**, GM-CSF average TAM expression in samples where proinflammatory macrophages are majority or not, as previously classified in the same cohort (3). Metastasizing and non-metastasizing patients, are also indicated. Scale bar, 50 μm.

### GM-CSF enhances invasion and metastasis of melanoma cells

As GM-CSFR subunits were high and widely expressed by melanoma cells in the majority of primary tumors, we explored whether the cytokine could directly affect to melanoma cell functions. We first confirmed the expression of both CD116 and CD131 in distinct melanoma cell lines by immunoblotting (Fig. 2A), and explored the distinct potential signaling pathways known to be triggered by different concentrations of GM-CSF. Differently to monocytes, where the cytokine signals through canonical JAK2/STAT5 pathway, melanoma cell lines exposed to exogenous human GM-CSF showed activation of STAT3, and to a lower extent PI3K/AKT, at the time and doses tested, demonstrating that the cytokine receptor is functional and may affect essential functions of melanoma cells (Fig. 2A and supplementary Fig. S1A and S1B). Thus, to determine the in vitro effects of GM-CSF, we performed tumor cell invasion and serum-deprived survival assays. BLM and Skmel-103 melanoma spheroids were embedded in collagen matrices in the absence or presence of exogenous GM-CSF, showing specific dose-dependent invasive responses that could be blocked with neutralizing anti-CD116 antibody, whereas no invasive response was observed for human M-CSF (Fig. 2B). Whereas invasion was enhanced by GM-CSF, survival was not affected by the exogenous administration of the cytokine in our setting conditions (Fig. 2C). As invasiveness and metastatic dissemination are usually coupled capabilities of melanoma cells, we explored the in vivo effect of the cytokine on the colonization process of a distant organ in a murine model. BLM and A375 melanoma cells were intravenously injected, together with human GM-CSF and anti-CD116 or control isotype antibodies, to mimic circulating TCs (Fig. 2D). Human GM-CSF, which acts exclusively on human cells (23), significantly heightened melanoma cells lung colonization, an effect that could be reversed with anti-CD116 blocking antibody (Fig. 2E and 2F). Altogether our data suggest that GM-CSF has a pro-tumoral and pro-metastatic direct effect on melanoma cells, which is in line with the enriched cytokine expression observed in metastasizing primary tissues.

**Figure 2.**
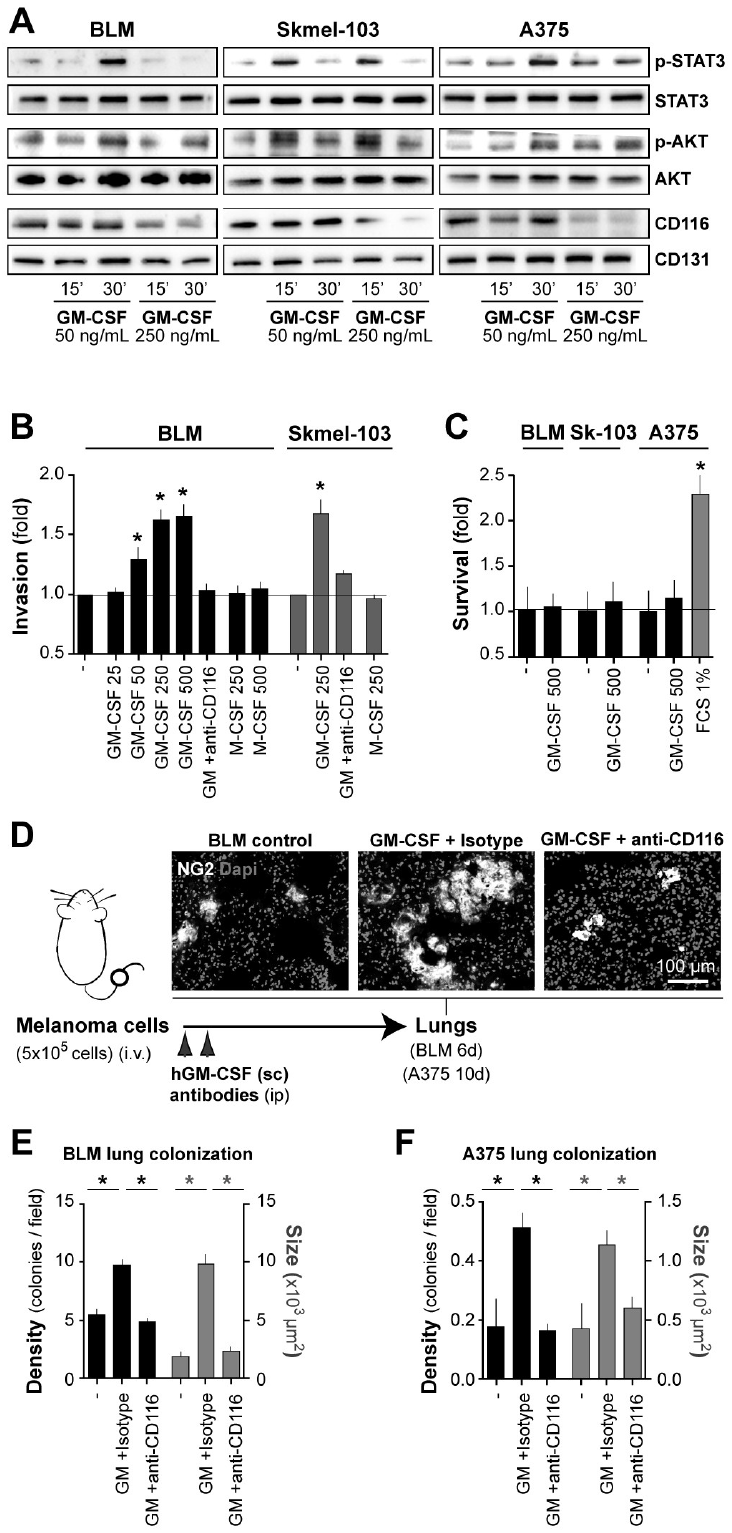
Direct effect of GM-CSF on melanoma cells. **A**, GM-CSF signaling kinetics in BLM, Skmel-103 and A375 melanoma cell lines whole lysates following stimulation with GM-CSF at 50 or 250 ng/mL for 0, 15, and 30 minutes. Immunoblots show phosphorylated STAT3 (Tyr705) and AKT (Thr308) and total forms of the proteins, alongside with CD116 and CD131. Blots are representative of 2-3 independent experiments. **B**, BLM and Skmel-103 spheroids embedded in 3D collagen and allowed to invade for 72 hours in the presence of increasing concentrations of exogenous GM-CSF (ng/ml), as indicated. Treatment with M-CSF was also included, as negative control. Functionality of GM-CSF receptors on melanoma cells was assessed using neutralizing antibodies against CD116 and its corresponding control isotype. Mean ±SD fold-change invasion values are shown (n= 3-6). **C**, 5-days serum-deprivation survival assays of melanoma cell lines in the absence or presence of 500 ng/mL GM-CSF. FCS 1% condition is included in Skmel-103, as positive control. Mean ±SD fold-change optical density values are shown (n= 3-5). **D**, scheme of lung colonization model assay in immune-compromised NSG mice. NG2^+^ BLM colonies are shown in representative lung sections from mice treated or not with exogenous human GM-CSF and neutralizing antibodies against CD116 or isotype control, as indicated. **E, F**, plots showing BLM and A375 lung colonies quantifications after 7 and 10 days of intravenous injection, respectively. Mean ± SD density (number/field) and size (μm^2^) values of spontaneous melanoma micrometastases are shown (n= 4-5 mice per condition). In all panels, unpaired *t*-test analyses are shown; *\*p*< 0.05 (relative to respective controls).

### GM-CSF-primed macrophages display enhanced activation and inflammatory cytokine secretion in response to melanoma cells

As monocytes are recruited into tumors, which may be locally enriched in CSFs, we designed an in vitro approach to explore the functional effects of priming monocytes with either none, M-CSF or GM-CSF cytokines, in co-culture with melanoma cells. Monocytes/macrophages and melanoma cells were then separated and lysed for transcriptomic analysis after 24-hours coculture, whereas conditioned media (CM) were collected after 48-hours for secretome analysis (Fig. 3A). GM-CSF primed macrophages (GM-MPs) co-cultured with melanoma cells showed in all cases the highest secretion levels of inflammatory cytokines, including GM-CSF itself and its downstream effector Activin A (Fig. 3B), as well as the pro-metastatic chemokine CCL20 and the classical pro-inflammatory mediator IL-1β (supplementary Fig. S2A). M-CSF monocyte priming (M-MPs) in co-culture with melanoma cells led to the opposite effect on GM-CSF, Activin A and IL-1β secretion, when compared with unprimed conditions (Fig. 3B and supplementary Fig. S2A). Accordingly, melanoma-activated GM-MPs exhibited significantly higher expression of tumor-promoting genes such as *CSF2, INHBA, IL1B, CCL20, VEGFA, TNFA, PTGS2* or *IDO1* (Fig. 3C and supplementary Fig. S2B), than melanoma-activated M-MPs or monocytes. Otherwise, *CSF1* mRNA expression was diminished by melanoma co-culture in all priming-conditions (supplementary Fig. S2B). As it will be addressed later, GM-MPs not only experienced the maximal activation by TCs, but they also reciprocally exerted it on melanoma cells, as shown by significantly enhanced TC levels of *CSF2, INHBA* and *IL1B* mRNA (Fig. 3D). In order to decipher the functional effect of these conditioned media, invasion assays with melanoma cells were performed. BLM and Skmel-103 spheroids embedded in collagen matrices were incubated with different CM and allowed to invade for 72 hours. Conditioned media from melanoma cells cocultured with GM-MPs induced the highest invasion levels, while those from TCs co-cultured with the other experimentally defined macrophage subsets slightly or did not promoted melanoma invasion (Fig. 3E). Altogether, our findings support a model in which GM-CSF-primed monocytes exposed to melanoma cells are reprogrammed towards a highly pro-inflammatory state with a pro-metastatic secretome, as opposed to M-CSF-primed or unprimed monocytes.

**Figure 3.**
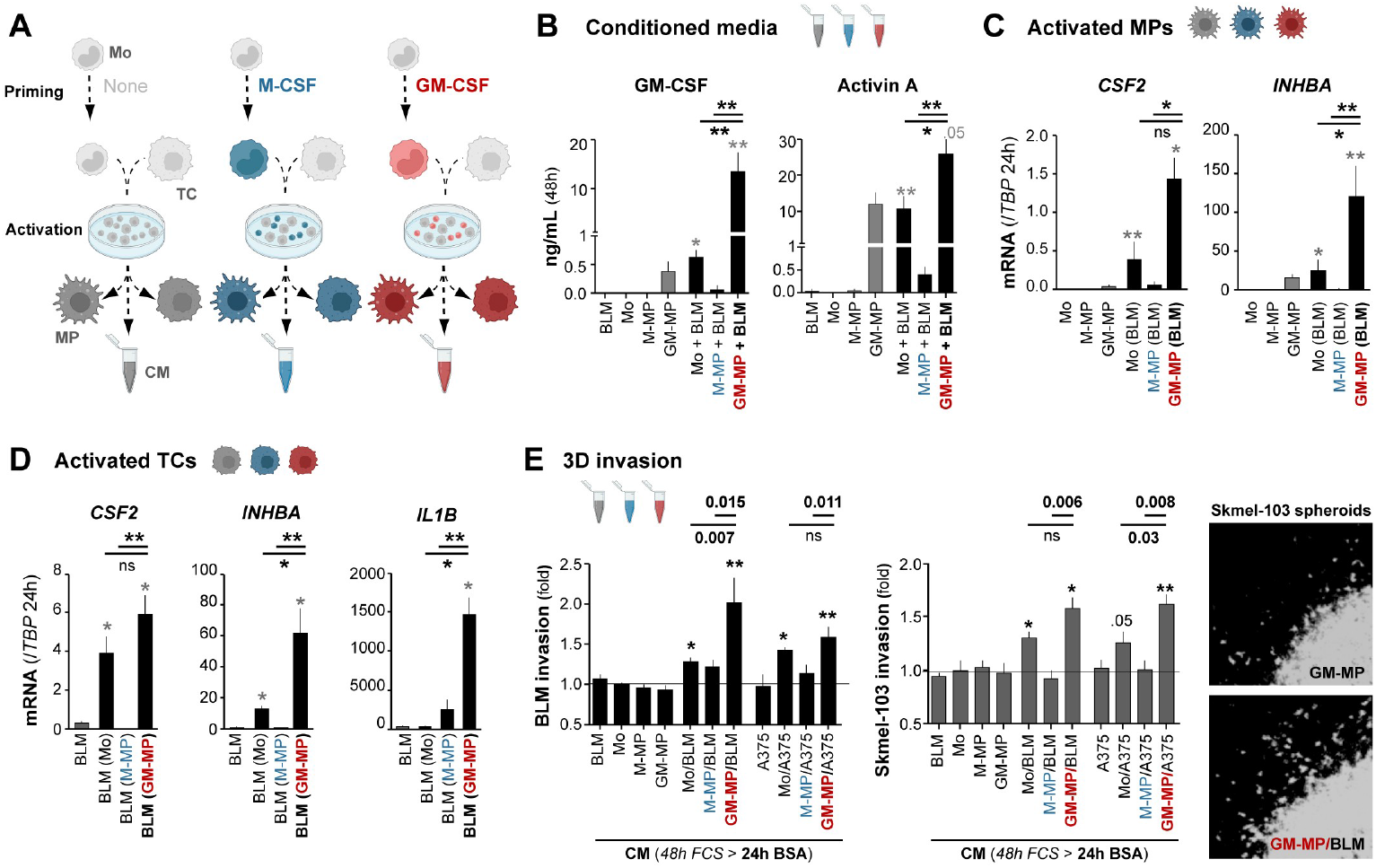
Protumoral loop induced by GM-CSF priming of monocytes and subsequent activation with melanoma cells. **A**, schematic representation of our model of monocyte priming with none, GM-CSF or M-CSF cytokines followed by co-culture with melanoma cells. All, conditioned media (CM) and mutually activated monocytes/macrophages and TCs, were collected or isolated for secretomic and transcriptomic analyses, respectively. **B**, secretion of GM-CSF and Activin A measured at 48-hours co-culture CM. Mean ±SD values are shown (n= 3-6 monocyte donors). **C, D**, *CSF2, INHBA* and/or *IL1B* mRNA expression quantified by qPCR in isolated monocytes/macrophages (**C**) or BLM TCs (**D**) after 24-hours co-culture. Mean ±SD values relative to *TBP* mRNA, are shown (n= 3-6 donors). **E**, BLM and Skmel-103 spheroids embedded in 3D-collagen and allowed to invade for 72 hours in the presence of indicated CM (see representative images of invading cells from the spheroid border). Mean ±SD fold-change invasion values are shown (n= 3-6). Paired *t*-test statistical values are shown or represented (relative to TCs-alone CM, *\*p* < 0.05; *\*\*p* < 0.005).

### GM-CSF-primed macrophages induce a profound transcriptomic reprogramming of melanoma cells

To better understand the reciprocal conditioning of monocytes/macrophages and melanoma cells, we performed RNA-sequencing to explore more deeply the transcriptome of activated melanoma cells. Differential expression analyses revealed that the most extensive and significant genetic changes occurred when BLM or A375 melanoma cells were co-cultured with GM-MPs, compared to M-MPs or unprimed monocytes. (Fig. 4A and supplementary Fig. S3A). Indeed, when directly comparing TCs conditioned by GM-MPs to those done by M-MPs or unprimed monocytes, we detected over a thousand of differentially expressed genes (DEGs) exclusively upregulated by GM-MPs that were primarily associated with inflammatory responses and the NF-κB signaling pathway (Fig. 4B and supplementary Fig. S3B). This inflammatory reprogramming was further confirmed by an enrichment analysis of cytokine production gene sets, where both BLM and A375 cells activated by GM-MPs showed maximal induction of pro-inflammatory cytokines, excluding type I IFNs (Fig. 4C and supplementary Fig. S3C). Altogether, these findings suggest that GM-MP reprogrammed melanoma cells, unlike those exposed to M-MPs or monocytes, exhibit a more active transcriptomic profile related to inflammatory responses, likely playing a key role in shaping its pro-invasive effect through secretome activation.

**Figure 4.**
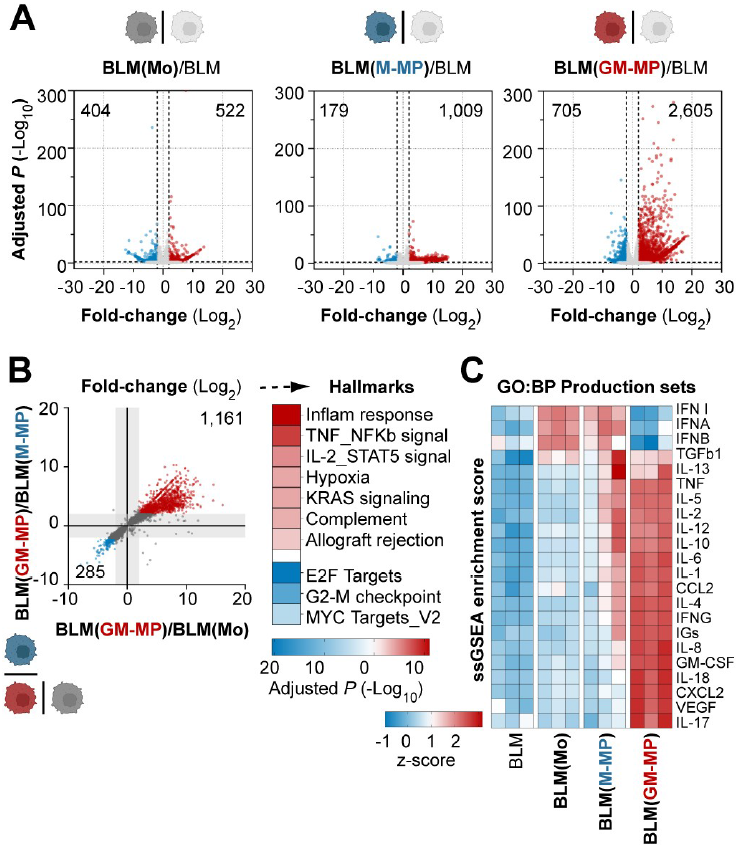
Differential gene expression and enrichment analyses of BLM cells activated by primed-macrophages. **A**, volcano plots showing differentially expressed genes (DEGs) for 24-hours melanoma cells co-cultured with monocytes (Mo) or primed-MPs vs unconditioned TCs, as indicated. Number of upregulated (red spots) or downregulated (blue) genes, are shown. **B**, DEG analysis of GM-MP-reprogrammed BLM cells compared to monocyte (*x*-axis) and M-MP (*y*-axis) activated BLM cells (Fold-change Log_2_> 2; Adjusted P< 0.05), and resultant hallmark enrichment analysis. Major significant gene sets (Adjusted P< 0.01), are shown. **C**, corresponding heatmap of single sample GSEA (ssGSEA) enrichment scores for selected gene sets related to production of cytokines from GO:BP (Gene Ontology, Biology Processes).

## DISCUSSION

As GM-CSF has been described to exert both immune-dependent and -independent functions, as well as anti-tumor and pro-tumoral effects across different cancer types, we first aimed to characterize its expression in primary melanoma tissues and evaluate its correlation with patient outcome. Remarkably, a clear inverse association exists between GM-CSF expression on both TAMs and melanoma cells and patient survival, becoming a novel independent prognostic marker in cutaneous melanoma. This pro-tumoral correlation aligns with previous breast and liver tumor studies (24,25), but differs from colorectal cancer studies where GM-CSF is indicative of favorable prognosis (26,27). In our study we have focused on the pro-tumoral role of GM-CSF and have explored the mechanisms operating in TAMs and melanoma cells. Thus, exogenous GM-CSF was shown to signal through alternative pathways STAT3 and AKT in melanoma cells, rather than the canonical JAK2/STAT5 pathway. These activated proteins are known to be involved in melanoma progression when aberrantly stimulated by different cytokines and growth factors (28,29), which now include GM-CSF in melanoma patients with a poor prognosis. Of note, GM-CSF released by several cancer cell types has been linked to PD-L1 expression in distinct myeloid cells through JAK2/STAT3 activation (9,30,31), suggesting that a similar auto/paracrine axis might be operative in melanoma cells from metastasizing tumors, where PD-L1 is also upregulated (3).

Beyond their role in the immune system, cytokines may exert a direct influence on cancerous cells, and GM-CSF has been suggested to promote cell proliferation and/or migration in a variety of solid tumors and cancer cell lines (14,32). Specifically, GM-CSF has been shown to induce proliferation of breast and brain cancer cells (33,34), invasion of colon cancer cells (35), or both functions in head and neck squamous cell carcinoma (HSNCC) cells (36). However, GM-CSF role in human melanoma is yet to be determined (11). Here we show how exogenous GM-CSF may directly enhance the invasive capacity of melanoma cells in vitro, rather than promoting their survival; and further confirm its pro-metastatic effect in a mouse lung colonization model. A mouse model with HSNCC had previously shown a direct pro-tumoral role of GM-CSF on tumor growth, but only when acting synergistically with G-CSF (13). Thus, the wide and constant expression of GM-CSFR on primary melanoma cells, together with the increased expression of GM-CSF in metastasizing tumors, suggest that an auto/paracrine loop with pro-tumoral effects may be established along melanoma evolution, eventually promoting distant metastasis.

The complex and controversial role of GM-CSF in cancer is also due to its indirect pleiotropic effects on the immune system, including macrophages, neutrophils, and dendritic cells. Specifically, TAMs can inhibit antitumor immune responses by secreting cytokines, chemokines, and growth factors, indirectly promoting tumor growth and progression (11). We have demonstrated that GM-CSF-primed monocytes are capable of reprogramming melanoma cells, promoting a pro-inflammatory cytokine microenvironment and enhancing tumor invasion. This pro-tumoral priming effect has been previously shown in other cancer cell types (37,38). Moreover, our study suggests that both GM-CSF-primed monocytes and melanoma cells acquire a pro-inflammatory secretory phenotype after co-culture. This effect was not observed in monocytes or M-MPs, reflecting a unique amplified response of macrophages and TCs crosstalk in GM-CSF rich microenvironments. Hence, as suggested by Osuala et al. in breast cancer, GM-CSF secretion by melanoma cells might act as a feed-forward loop, priming myeloid cells to produce pro-inflammatory cytokines and growth factors that may drive tumor progression (39). Our prior work showing that pro-inflammatory cytokine-producing TAMs are associated with poor prognosis in human melanoma, would support this model (3).

Exogenous GM-CSF has been explored as an adjuvant to distinct cancer immunotherapies, but its value remains controversial due to its variable effects on tumor progression (14,40). The previously unnoticed GM-CSF auto/paracrine axis occurring between TAMs and TCs might help explain, at least partially, the lack of therapeutic effect of the cytokine when administered to patients with resected stage III-IV melanomas (41–43). Indeed, a randomized trial was conducted in melanoma patients to compare the effects of subcutaneous GM-CSF versus intralesional Talimogene Laherparepvec (T-VEC), a herpes simplex virus genetically modified to produce GM-CSF locally. Although T-VEC showed improved durable response rates in patients with advanced melanoma, it did not improve overall survival (44), and the specific role of GM-CSF in this response was not addressed. To achieve a beneficial effect in patients, GM-CSF is likely needed to be combined with other therapeutic agents capable of concomitantly modifying the tumor microenvironment (45). In this regard, a recent preclinical study has shown that GM-CSF and IL-12 cytokines, modified to be retained at the tumoral stroma, synergistically enhance TAM and CD8^+^ cell populations against B16 melanoma tumors in mice (46).

On the contrary, other preclinical mouse models of pancreatic cancer have shown that neutralization of tumor-derived GM-CSF effectively impedes tumor development (47), and depletion of GM-CSF in mesenchymal cells inhibits the ability of these cells to promote tumor growth and metastasis (48). Therefore, understanding the signaling networks existing in the melanoma microenvironment may contribute to the development of efficient targeted therapies to halt tumor progression and/or metastasis. In this case, the existence of a GM-CSF auto/paracrine axis in melanoma patients with poor-prognosis points to blockade of the axis, rather than supporting it, as a therapeutic avenue worth exploring.

## Acknowledgments

We thank Alejandra de Francisco, José María Bellón and the animal facility staff for their expert technical assistance. This work was supported by the Ministry of Science and Innovation PID2021-123507OB-I00 grant (PSM, RS) co-financed by ERDF/FEDER Funds from the European Commission, ‘A way of making Europe’; and by the Accelerator Award Program C18915/A29362 grant and the Dirección General de Innovación e Investigación Tecnológica de la Comunidad de Madrid P2022/BMD-7274 grant. AN-V, CB-A and AG-S were financed by the Comunidad de Madrid YEI program.

## Data availability statement

The data generated in this study are publicly available in Gene Expression Omnibus (GEO) at GSE171277, GSE242674 and GSE280027.

## Author contributions

Conceptualization: EG-M, PS-M, RS; Data Curation: AN-V, CB-A, AG-S, JAA-I, RS; Formal Analysis: EG-M, AN-V, CB-A, BL-N, AG-S, RS; Funding acquisition: PS-M, RS; Investigation: EG-M, AN-V, CB-A, BL-N, AG-S, RS; Methodology: EG-M, AN-V, CB-A, BL-N, RS; Project administration: EG-M, RS; Resources: JAA-I, VP-B, PS-M, RS; Supervision: RS; Validation: EG-M, AN-V, CB-A, RS; Writing-Original Draft Preparation: EG-M, RS; Writing-Review and Editing: AN-V, CB-A, BL-N, P-SM.

## LEGENDS

**Supplementary Figure 1. A, B**, GM-CSF signaling kinetics in BLM, Skmel-103 and A375 melanoma cell lines (**A**), as well as fresh-monocyte whole lysates (**B**), following stimulation with indicated GM-CSF doses for 15 and/or 30 minutes. Control lines in A and B correspond to 30 and 15 minutes untreated cells, respectively. Immunoblots show phosphorylated STAT3 (Tyr705), AKT (Thr308), p65 (Ser536), Erk (Thr202/Tyr204), STAT1 (Tyr701), and/or STAT5 (Tyr694/699), and their respective total form of the proteins. Blots are representative of 2-3 independent experiments.

**Supplementary Figure 2. A**, CCL20 and IL-1β secretion assessed at CM of monocytes/primed-macrophages cocultured or not for 48 hours with BLM cells, as indicated. Mean ±SD values are shown (n= 3-6 donors). **B**, mRNA expression quantified by qPCR in isolated monocytes or primed-MPs after 24-hours co-culture with BLM cells, as indicated. Mean ± SD values relative to *TBP* mRNA, are shown (n = 3-6 donors).

**Supplementary Figure 3. A**, volcano plots showing DEGs for 24-hours A375 cells activated by monocytes or primed-MPs vs unconditioned A375 cells, as indicated. Number of upregulated (red spots) or downregulated (blue) genes, are shown. **B**, differential gene expression analysis of GM-MP-reprogrammed A375 cells compared to Mo-activated (*x*-axis) and M-MP-activated (*y*-axis) A375 cells (fold-change, Log_2_), and hallmark enrichment analysis of resultant upregulated (red spots) and downregulated (blue spots) genes. Major significant gene sets (Adjusted P< 0.01), are shown. **C**, corresponding heatmap of single sample GSEA (ssGSEA) enrichment scores for selected gene sets related to production of cytokines from GO:BP (Gene Ontology, Biology Processes).

**Supplementary Table 1**. Oligonucleotides and antibodies used in this study.

**Supplementary Table 2**. Mann-Whitney correlation analyses between clinico-pathologic features and GM-CSF expression.

## REFERENCES

1. Ma R-Y, Black A, Qian B-Z. Macrophage diversity in cancer revisited in the era of single-cell omics. Trends in Immunology. Elsevier; 2022;43:546–63.

2. Mulder K, Patel AA, Kong WT, Piot C, Halitzki E, Dunsmore G, et al. Cross-tissue single-cell landscape of human monocytes and macrophages in health and disease. Immunity. Elsevier; 2021;54:1883-1900.e5.

3. Barrio-Alonso C, Nieto-Valle A, Barandalla-Revilla L, Avilés-Izquierdo JA, Parra-Blanco V, Sánchez-Mateos P, et al. Translating genetics into tissue: inflammatory cytokine-producing TAMs and PD-L1 tumor expression as poor prognosis factors in cutaneous melanoma. Frontiers in Immunology. 2025;16:2025.03.04.640087.

4. Jeannin P, Paolini L, Adam C, Delneste Y. The roles of CSFs on the functional polarization of tumor-associated macrophages. The FEBS Journal. 2018;285:680–99.

5. Rodriguez RM, Suarez-Alvarez B, Lavín JL, Ascensión AM, Gonzalez M, Lozano JJ, et al. Signal Integration and Transcriptional Regulation of the Inflammatory Response Mediated by the GM-/M-CSF Signaling Axis in Human Monocytes. Cell Reports. Elsevier; 2019;29:860-872.e5.

6. Bhattacharya P, Budnick I, Singh M, Thiruppathi M, Alharshawi K, Elshabrawy H, et al. Dual Role of GM-CSF as a Pro-Inflammatory and a Regulatory Cytokine: Implications for Immune Therapy. J Interferon Cytokine Res. 2015;35:585–99.

7. Hercus TR, Broughton Sophie E., Ekert Paul G., Ramshaw Hayley S., Perugini Michelle, Grimbaldeston, Michele, et al. The GM-CSF receptor family: Mechanism of activation and implications for disease. Growth Factors. Taylor & Francis; 2012;30:63–75.

8. Guerrero P, Bono C, Sobén M, Guiu A, Cheng QJ, Gil ML, et al. GM-CSF receptor expression determines opposing innate memory phenotypes at different stages of myelopoiesis. Blood. 2024;143:2763–77.

9. Wang P, Tao L, Yu Y, Wang Q, Ye P, Sun Y, et al. Oral squamous cell carcinoma cell-derived GM-CSF regulates PD-L1 expression in tumor-associated macrophages through the JAK2/STAT3 signaling pathway. Am J Cancer Res. 2023;13:589–601.

10. Su X, Xu Y, Fox GC, Xiang J, Kwakwa KA, Davis JL, et al. Breast cancer–derived GM-CSF regulates arginase 1 in myeloid cells to promote an immunosuppressive microenvironment. J Clin Invest. 131:e145296.

11. Kumar A, Taghi Khani A, Sanchez Ortiz A, Swaminathan S. GM-CSF: A Double-Edged Sword in Cancer Immunotherapy. Front Immunol. 2022;13:901277.

12. Lacey DC, Achuthan A, Fleetwood AJ, Dinh H, Roiniotis J, Scholz GM, et al. Defining GM-CSF– and Macrophage-CSF–Dependent Macrophage Responses by In Vitro Models. The Journal of Immunology. 2012;188:5752–65.

13. Obermueller E, Vosseler S, Fusenig NE, Mueller MM. Cooperative autocrine and paracrine functions of granulocyte colony-stimulating factor and granulocyte-macrophage colony-stimulating factor in the progression of skin carcinoma cells. Cancer Res. 2004;64:7801– 12.

14. Hong I-S. Stimulatory versus suppressive effects of GM-CSF on tumor progression in multiple cancer types. Exp Mol Med. 2016;48:e242.

15. Zheng Q, Li X, Cheng X, Cui T, Zhuo Y, Ma W, et al. Granulocyte-macrophage colony-stimulating factor increases tumor growth and angiogenesis directly by promoting endothelial cell function and indirectly by enhancing the mobilization and recruitment of proangiogenic granulocytes. Tumour Biol. SAGE Publications Ltd STM; 2017;39:1010428317692232.

16. Roda JM, Wang Y, Sumner LA, Phillips GS, Marsh CB, Eubank TD. Stabilization of HIF-2α induces sVEGFR-1 production from tumor-associated macrophages and decreases tumor growth in a murine melanoma model. J Immunol. 2012;189:3168–77.

17. Moshe A, Izraely S, Sagi-Assif O, Malka S, Ben-Menachem S, Meshel T, et al. Inter-Tumor Heterogeneity—Melanomas Respond Differently to GM-CSF-Mediated Activation. Cells. 2020;9:1683.

18. Mehta A, Motavaf M, Nebo I, Luyten S, Osei-Opare KD, Gru AA. Advancements in Melanoma Treatment: A Review of PD-1 Inhibitors, T-VEC, mRNA Vaccines, and Tumor-Infiltrating Lymphocyte Therapy in an Evolving Landscape of Immunotherapy. J Clin Med. 2025;14:1200.

19. Gutiérrez-Seijo A, García-Martínez E, Barrio-Alonso C, Pareja-Malagón M, Acosta-Ocampo A, Fernández-Santos ME, et al. CCL20/TNF/VEGFA Cytokine Secretory Phenotype of Tumor-Associated Macrophages Is a Negative Prognostic Factor in Cutaneous Melanoma. Cancers (Basel). 2021;13:3943.

20. Barrio-Alonso C, Nieto-Valle A, García-Martínez E, Gutiérrez-Seijo A, Parra-Blanco V, Márquez-Rodas I, et al. Chemokine profiling of melanoma-macrophage crosstalk identifies CCL8 and CCL15 as prognostic factors in cutaneous melanoma. J Pathol. 2024;262:495– 504.

21. Gutiérrez-Seijo A, García-Martínez E, Barrio-Alonso C, Parra-Blanco V, Avilés-Izquierdo JA, Sánchez-Mateos P, et al. Activin A Sustains the Metastatic Phenotype of Tumor-Associated Macrophages and Is a Prognostic Marker in Human Cutaneous Melanoma. J Invest Dermatol. 2022;142:653-661.e2.

22. Wu T, Hu E, Xu S, Chen M, Guo P, Dai Z, et al. clusterProfiler 4.0: A universal enrichment tool for interpreting omics data. Innovation (Camb). 2021;2:100141.

23. Nishijima I, Nakahata T, Hirabayashi Y, Inoue T, Kurata H, Miyajima A, et al. A human GM-CSF receptor expressed in transgenic mice stimulates proliferation and differentiation of hemopoietic progenitors to all lineages in response to human GM-CSF. Mol Biol Cell. 1995;6:497–508.

24. Su S, Liu Q, Chen J, Chen J, Chen F, He C, et al. A positive feedback loop between mesenchymal-like cancer cells and macrophages is essential to breast cancer metastasis. Cancer Cell. 2014;25:605–20.

25. Ruffolo LI, Jackson KM, Kuhlers PC, Dale BS, Figueroa Guilliani NM, Ullman NA, et al. GM-CSF drives myelopoiesis, recruitment and polarisation of tumour-associated macrophages in cholangiocarcinoma and systemic blockade facilitates antitumour immunity. Gut. 2022;71:1386–98.

26. Urdinguio RG, Fernandez AF, Moncada-Pazos A, Huidobro C, Rodriguez RM, Ferrero C, et al. Immune-dependent and independent antitumor activity of GM-CSF aberrantly expressed by mouse and human colorectal tumors. Cancer Res. 2013;73:395–405.

27. Nebiker CA, Han J, Eppenberger-Castori S, Iezzi G, Hirt C, Amicarella F, et al. GM-CSF Production by Tumor Cells Is Associated with Improved Survival in Colorectal Cancer. Clin Cancer Res. 2014;20:3094–106.

28. Dai DL, Martinka M, Li G. Prognostic significance of activated Akt expression in melanoma: a clinicopathologic study of 292 cases. J Clin Oncol. 2005;23:1473–82.

29. Logotheti S, Pützer BM. STAT3 and STAT5 Targeting for Simultaneous Management of Melanoma and Autoimmune Diseases. Cancers (Basel). 2019;11:1448.

30. Wang T-T, Zhao Y-L, Peng L-S, Chen N, Chen W, Lv Y-P, et al. Tumour-activated neutrophils in gastric cancer foster immune suppression and disease progression through GM-CSF-PD-L1 pathway. Gut. 2017;66:1900–11.

31. Thorn M, Guha P, Cunetta M, Espat NJ, Miller G, Junghans RP, et al. Tumor-associated GM-CSF overexpression induces immunoinhibitory molecules via STAT3 in myeloid-suppressor cells infiltrating liver metastases. Cancer Gene Ther. 2016;23:188–98.

32. Aliper AM, Frieden-Korovkina VP, Buzdin A, Roumiantsev SA, Zhavoronkov A. A role for G-CSF and GM-CSF in nonmyeloid cancers. Cancer Med. 2014;3:737–46.

33. Ansari KI, Bhan A, Saotome M, Tyagi A, De Kumar B, Chen C, et al. Autocrine GMCSF Signaling Contributes to Growth of HER2+ Breast Leptomeningeal Carcinomatosis. Cancer Res. 2021;81:4723–35.

34. Revoltella RP, Menicagli M, Campani D. Granulocyte-macrophage colony-stimulating factor as an autocrine survival-growth factor in human gliomas. Cytokine. 2012;57:347–59.

35. Gutschalk CM, Yanamandra AK, Linde N, Meides A, Depner S, Mueller MM. GM-CSF enhances tumor invasion by elevated MMP-2, -9, and -26 expression. Cancer Med. 2013;2:117–29.

36. Gutschalk CM, Herold-Mende CC, Fusenig NE, Mueller MM. Granulocyte colony-stimulating factor and granulocyte-macrophage colony-stimulating factor promote malignant growth of cells from head and neck squamous cell carcinomas in vivo. Cancer Res. 2006;66:8026–36.

37. Huang D, Song S-J, Wu Z-Z, Wu W, Cui X-Y, Chen J-N, et al. Epstein-Barr Virus-Induced VEGF and GM-CSF Drive Nasopharyngeal Carcinoma Metastasis via Recruitment and Activation of Macrophages. Cancer Res. 2017;77:3591–604.

38. Rigo A, Gottardi M, Zamò A, Mauri P, Bonifacio M, Krampera M, et al. Macrophages may promote cancer growth via a GM-CSF/HB-EGF paracrine loop that is enhanced by CXCL12. Mol Cancer. 2010;9:273.

39. Osuala KO, Chalasani A, Aggarwal N, Ji K, Moin K. Paracrine Activation of STAT3 Drives GM-CSF Expression in Breast Carcinoma Cells, Generating a Symbiotic Signaling Network with Breast Carcinoma-Associated Fibroblasts. Cancers (Basel). 2024;16:2910.

40. Mihalik NE, Steinberger KJ, Stevens AM, Bobko AA, Hoblitzell EH, Tseytlin O, et al. Dose-Specific Intratumoral GM-CSF Modulates Breast Tumor Oxygenation and Antitumor Immunity. J Immunol. 2023;211:1589–604.

41. Faries MB, Hsueh EC, Ye X, Hoban M, Morton DL. Effect of granulocyte/macrophage colony-stimulating factor on vaccination with an allogeneic whole-cell melanoma vaccine. Clin Cancer Res. 2009;15:7029–35.

42. Hodi FS, Lee S, McDermott DF, Rao UN, Butterfield LH, Tarhini AA, et al. Ipilimumab plus sargramostim vs ipilimumab alone for treatment of metastatic melanoma: a randomized clinical trial. JAMA. 2014;312:1744–53.

43. Lawson DH, Lee S, Zhao F, Tarhini AA, Margolin KA, Ernstoff MS, et al. Randomized, Placebo-Controlled, Phase III Trial of Yeast-Derived Granulocyte-Macrophage Colony-Stimulating Factor (GM-CSF) Versus Peptide Vaccination Versus GM-CSF Plus Peptide Vaccination Versus Placebo in Patients With No Evidence of Disease After Complete Surgical Resection of Locally Advanced and/or Stage IV Melanoma: A Trial of the Eastern Cooperative Oncology Group-American College of Radiology Imaging Network Cancer Research Group (E4697). J Clin Oncol. 2015;33:4066–76.

44. Andtbacka RHI, Collichio F, Harrington KJ, Middleton MR, Downey G, Öhrling K, et al. Final analyses of OPTiM: a randomized phase III trial of talimogene laherparepvec versus granulocyte-macrophage colony-stimulating factor in unresectable stage III-IV melanoma. J Immunother Cancer. 2019;7:145.

45. Ribas A, Dummer R, Puzanov I, VanderWalde A, Andtbacka RHI, Michielin O, et al. Oncolytic Virotherapy Promotes Intratumoral T Cell Infiltration and Improves Anti-PD-1 Immunotherapy. Cell. 2017;170:1109-1119.e10.

46. Kang S, Mansurov A, Kurtanich T, Chun HR, Slezak AJ, Volpatti LR, et al. Engineered GM-CSF polarizes protumorigenic tumor-associated macrophages to an antitumorigenic phenotype and potently synergizes with IL-12 immunotherapy. J Immunother Cancer. 2024;12:e009541.

47. Bayne LJ, Beatty GL, Jhala N, Clark CE, Rhim AD, Stanger BZ, et al. Tumor-derived granulocyte-macrophage colony-stimulating factor regulates myeloid inflammation and T cell immunity in pancreatic cancer. Cancer Cell. 2012;21:822–35.

48. Waghray M, Yalamanchili M, Dziubinski M, Zeinali M, Erkkinen M, Yang H, et al. GM-CSF Mediates Mesenchymal–Epithelial Cross-talk in Pancreatic Cancer. Cancer Discov. 2016;6:886–99.

