## Supplementary Table 1 for "Adverse prognosis of GM-CSF expression in human cutaneous melanoma"

| Gene | Primer Sequences (Roche) | Technique |
|---|---|---|
| *INHBA* | S: 5' ctcggagatcatcacgtttg; AS: 5' ccttggaaatctcgaagtgc | qPCR |
| *CSF2* | S: 5' tctcagaaatgtttgacctcca; AS: 5' gcccttgagcttggtgag | qPCR |
| *CCL20* | S: 5' gctgctttgatgtcagtgct; AS: 5' gcagtcaaagttgcttgctg | qPCR |
| *VEGFA* | S: 5' gcagcttgagttaaacgaacg; AS: 5' ggttcccgaaaccctgag | qPCR |
| *CSF1* | S: 5' gaactgccagtgtagagggaat; AS: 5' gctggtcagacaacatctgg | qPCR |
| *PTGS2* | S: 5' cttcacgcatcagtttttcaag; AS: 5' tcaccgtaaatatgatttaagtccac | qPCR |
| *IDO1* | S: 5' gtgtttcaccaaatccacga; AS: 5' ctgatagctgggggttgc | qPCR |
| *TNFA* | S: 5' cagcctcttctccttcctgat; AS: 5' gccagagggctgattagaga | qPCR |
| *IL1B* | S: 5' ctgtcctgcgtgttgaaaga; AS: 5' ttgggtaatttttgggatctaca | qPCR |

| Antigen | Antibody information | Technique (IF, immunofluorescence) |
|---|---|---|
| CD68 | Dako (Clone PG-M1), mouse IgG3 | IF-paraffin (pH 9.0) |
| GM-CSF | R&D MAB215, mouse IgG1 | IF-paraffin (pH 9.0) |
| GM-CSF | R&D AF215, Polyclonal Goat | IF-paraffin (pH 9.0) |
| M-CSF | Santa Cruz Biotechnology, sc-365779, mouse IgG2b | IF-paraffin (pH 9.0) |
| CD131 (GM-CSFR common β subunit) | R&D AF906, Polyclonal Goat | IF-paraffin (pH 9.0) |
| CD131 (GM-CSFR common β subunit) | Invitrogen MA5-48430, mouse IgG1 | Immunoblotting |
| CD116 (GM-CSFRα) | Santa Cruz Biotechnology, sc-456, mouse IgG1 | IF-paraffin (pH 9.0), Immunoblotting |
| CD116 (GM-CSFRα) | Mavrilimumab (CAM 3001), MedChemExpress, HY-P99031, human IgG4 | Neutralization assays |
| Phospho-STAT3 (Tyr705) | Cell Signaling, clone D3A7, rabbit | Immunoblotting |
| STAT3 | Cell Signaling, clone 79D7, rabbit | Immunoblotting |
| Phospho-AKT (Thr308) | Cell Signaling, #9275, rabbit | Immunoblotting |
| Akt (pan) | Cell Signaling, clone C67E7, rabbit | Immunoblotting |
| Phospho-NF-κB p65 (Ser536) | Cell Signaling, clone 93H1, rabbit | Immunoblotting |
| NF-κB p65 | Cell Signaling, clone D14E12, rabbit | Immunoblotting |
| Phospho-STAT1 (Tyr701) | Cell Signaling, clone D4A7, rabbit | Immunoblotting |
| STAT1 | Cell Signaling, #9172, rabbit | Immunoblotting |
| Phospho-STAT5A/B (Tyr694/699) | Millipore, clone 8-9-2 mouse | Immunoblotting |
| STAT5 | Cell Signaling, clone D2O6Y, rabbit | Immunoblotting |
| Phospho-ERK1/2 (Thr202/Tyr204) | Cell Signaling, clone D13.14.4E, rabbit | Immunoblotting |
| ERK | Cell Signaling, clone 137F5, rabbit | Immunoblotting |
| NG2 | BD Bioscience, 554275, mouse IgG2a | IF-frozen |
