## Supplementary Table 2 for "Adverse prognosis of GM-CSF expression in human cutaneous melanoma"

Sup TABLE 2 MANN-WHITNEY

| | | TAM GM-CSF | P | TC GM-CSF | P |
|---|---|---|---|---|---|
| **Sex** | Male (n=43) | 79 ± 5 | 0,493 | 67 ± 5 | 0.412 |
| | Female (n=37) | 85 ± 6 | | 75 ± 5 | |
| **Age** (years) | ≤65 (n=37) | 80 ± 5 | 0.806 | 74 ± 5 | 0.524 |
| | >65 (n=43) | 83 ± 6 | | 68 ± 5 | |
| **Location** | Head/limbs (n=52) | 86 ± 5 | 0.099 | 75 ± 5 | 0.138 |
| | Trunk (n=28) | 73 ± 5 | | 63 ± 6 | |
| **Subtype** | Nodular (n=30) | 80 ± 7 | 0.766 | 64 ± 6 | 0.257 |
| | Others (n=50) | 83 ± 5 | | 74 ± 4 | |
| **Ulceration** | No (n=39) | 84 ± 6 | 0.530 | 66 ± 5 | 0.313 |
| | Yes (n=37) | 78 ± 6 | | 74 ± 5 | |
| **Breslow** | 2-4 mm (n=41) | 87 ± 5 | 0.120 | 73 ± 5 | 0.405 |
| | >4 mm (n=39) | 75 ± 6 | | 68 ± 5 | |
| **Stage** | II (n=59) | 80 ± 5 | 0.544 | 67 ± 4 | 0.083 |
| | III-IV (n=21) | 84 ± 5 | | 81 ± 8 | |
| **Metastasis** (10 years) | No (n=35) | 58 ± 4 | <0.0001 | 50 ± 4 | <0.0001 |
| | Yes (n=45) | 99 ± 5 | | 87 ± 4 | |
| **Survival** (10 years) | Yes (n=50) | 78 ± 5 | 0.190 | 63 ± 4 | 0.002 |
| | No (n=30) | 87 ± 5 | | 81 ± 4 | |
