## Supplementary figures and images for "Adverse prognosis of GM-CSF expression in human cutaneous melanoma"

### Supplementary Figure 1

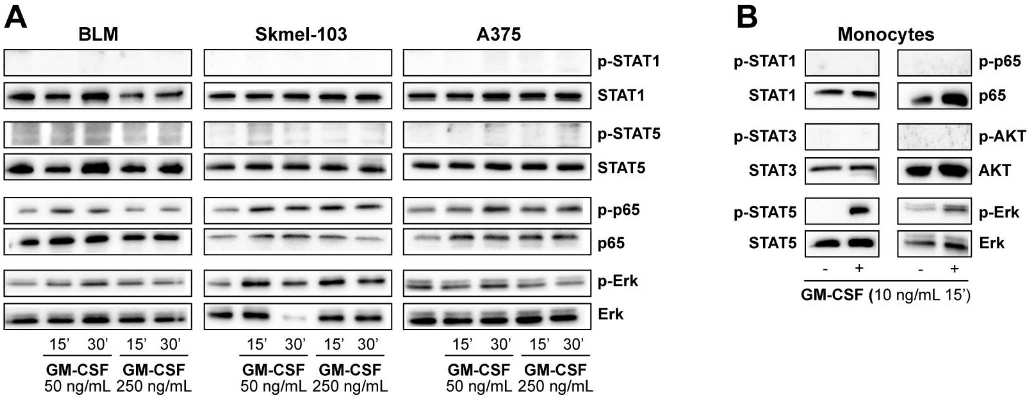


**Supplementary Figure 1**

### Supplementary Figure 2

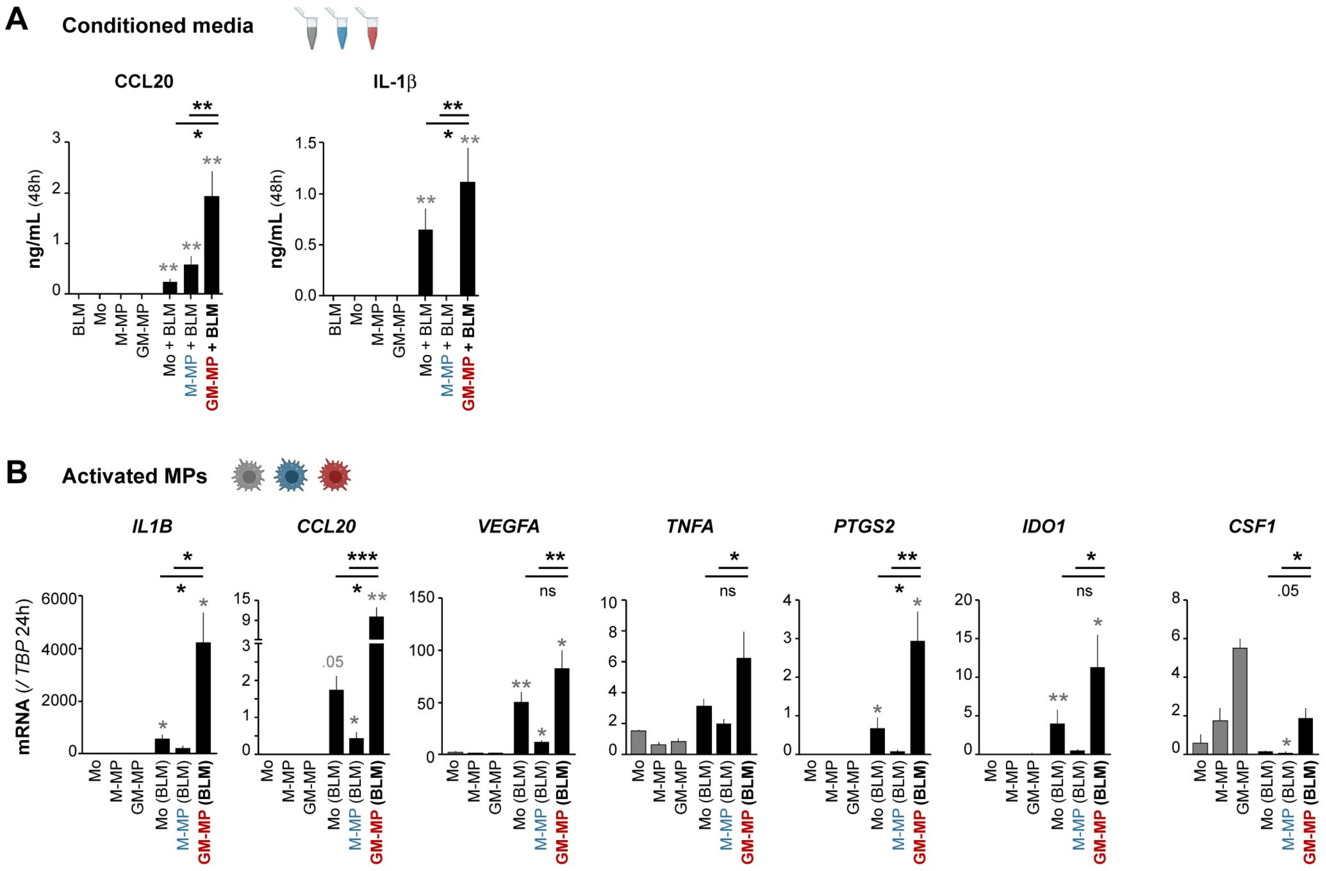


**Supplementary Figure 2**

### Supplementary Figure 3

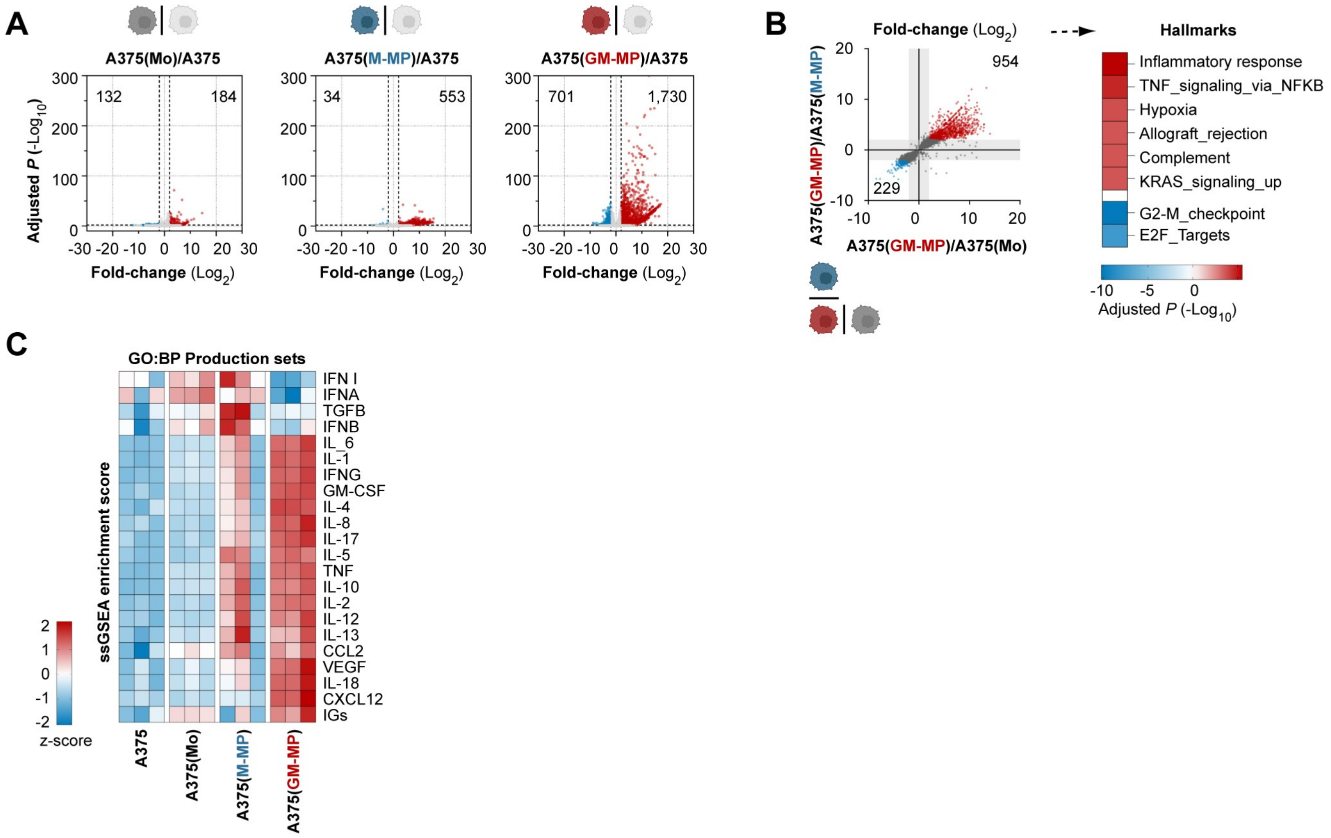


**Supplementary Figure 3**
